# Regulatory sites for known and novel splicing in human basal ganglia are enriched for disease-relevant information

**DOI:** 10.1101/591156

**Authors:** Sebastian Guelfi, Karishma D’Sa, Juan Botía, Jana Vandrovcova, Regina H. Reynolds, David Zhang, Daniah Trabzuni, Leonoardo Collado-Torres, Andrew Thomason, Pedro Quijada Leyton, Sarah A. Gagliano, Mike A. Nalls, UK Brain Expression Consortium, Kerrin S. Small, Colin Smith, Adaikalavan Ramasamy, John Hardy, Michael E. Weale, Mina Ryten

## Abstract

Genome-wide association studies have generated an increasing number of common genetic variants that affect neurological and psychiatric disease risk. Given that many causal variants are likely to operate by regulating gene expression, an improved understanding of the genetic control of gene expression in human brain is vital. However, the difficulties of sampling human brain, and its complexity, has meant that brain-related expression quantitative trait loci (eQTL) and allele specific expression (ASE) signals have been more limited in their explanatory power than might otherwise be expected. To address this, we use paired genomic and transcriptomic data from putamen and substantia nigra dissected from 117 brains, combined with a comprehensive set of analyses, to interrogate regulation at different stages of RNA processing and uncover novel transcripts. We identify disease-relevant regulatory loci and reveal the types of analyses and regulatory positions yielding the most disease-specific information. We find that splicing eQTLs are enriched for neuron-specific regulatory information; that ASE analyses provide highly cell-specific regulatory information; and that incomplete annotation of the brain transcriptome limits the interpretation of risk loci for neuropsychiatric disease. We release this rich resource of regulatory data through a searchable webserver, http://braineacv2.inf.um.es/.

## Introduction

The use of genome-wide genotyping in large patient and control populations has resulted in the identification of increasing numbers of common variants that impact on the risk of a wide range of neurological and psychiatric conditions, including Parkinson’s disease^1–5^, Alzheimer’s disease^6–10^ and schizophrenia^11,12^. However, the majority of these risk loci are still poorly characterised, and we do not yet fully understand the underlying molecular and cellular processes through which they act. As it is reasonable to assume that many causal variants operate by regulating gene expression, several studies have attempted to address this problem through the use of expression quantitative trait loci (eQTL) and allele-specific expression (ASE) analyses in a wide range of human tissues, with the aim of finding eQTL and ASE signals that colocalise with disease risk signals^13,14^.

This approach has had success, but perhaps not as much as might have been expected for all diseases. While the identification of eQTLs in blood have provided insights into autoimmune disorders^15–17^, the utility of brain-related eQTL and ASE data sets, particularly with regard to neurodegenerative disorders has been harder to demonstrate. For example, monocyte eQTL data sets appear to provide greater insights for Alzheimer’s disease^17^ probably because they reflect regulatory processes in microglia. This would suggest that while relevant eQTL and ASE signals are present in the brain, they are currently difficult to detect given the constraints on eQTL and ASE analyses in human brain.

Currently, the most easily available sampling method for brain tissue is post-mortem, making repeat sampling impossible and typically leading to smaller sample sizes, particularly for smaller structures such as the substantia nigra. Furthermore, the brain is a highly complex organ. Not only is it split into many regions with known inter-regional differences in expression^18^, each region is composed of an assemblage of different cell types, which complicates the interpretation of transcriptomic data and limits statistical power. Finally, the brain transcriptome is unusual in having a high degree of alternative splicing and a high degree of non-coding RNA activity^19^, much of which has yet to be fully characterised^20^.

We have addressed the latter of these constraints by conducting total RNA sequencing in two basal ganglia regions of clinical interest to human neurodegenerative and neuropsychiatric disorders: the substantia nigra and putamen. Using a comprehensive set of analyses to interrogate different stages of RNA processing and uncover novel unannotated transcripts, we have sought to identify not only disease-relevant regulatory loci but also the types of analyses and regulatory positions yielding the most brain and disease-specific information. We find that splicing eQTLs are enriched for neuron-specific regulatory information; that ASE analyses, probably by more effectively controlling for cellular heterogeneity, provide highly cell-specific regulatory information; and that incomplete annotation of the brain transcriptome is limiting the interpretation of risk loci for neuropsychiatric disease. We have released the rich resource of eQTL and ASE data generated in this study through a searchable webserver, http://braineacv2.inf.um.es/ (**Supplementary Figure 1**).

## Results

### RNA quantification and eQTL discovery

We assayed DNA and RNA from 180 brain samples originating from 117 individuals of European descent, which were part of the UK Brain Expression Consortium data set^21^ and which were classified as neurologically healthy based on the absence of neurological disease during life and neuropathological assessment (**Supplementary Table 1**). Using these paired data, we searched for eQTL associations between ~6.5 million genetic variants and ~411,000 RNA expression traits in putamen and ~370,000 RNA expression traits in substantia nigra, resulting in ~5.3 billion eQTL tests. We generated RNA expression traits using both annotation-based (known transcripts) and annotation-agnostic approaches (**Figure 1a**). Within annotated regions, RNA quantification was performed with RNA processing in mind (**Figure 1a**) to produce four separate measures of transcription, which formed the bases of our eQTL analysis. This resulted in the generation of four types of eQTLs (gi-eQTLs, e-eQTLs, ex-ex-eQTLs and ge-eQTLs) of which two were designed to capture the genetic regulation of splicing eQTLs (e-eQTLs and ex-ex-eQTLs). Finally, we included annotation-independent approaches to quantify transcription. We focused specifically on unannotated transcription within intergenic regions (producing i-eQTLs, **Online Methods**).

**Figure 1.**
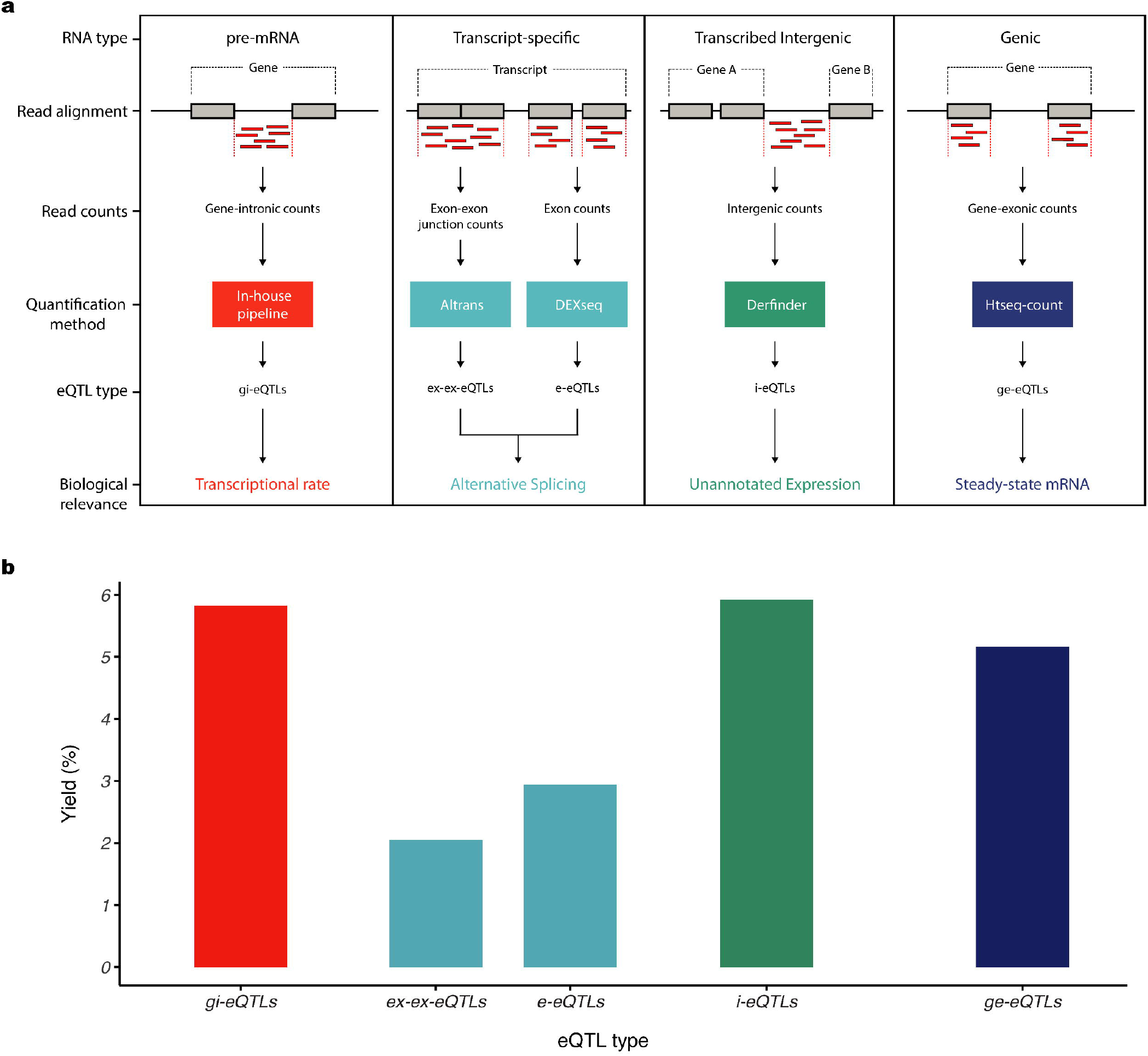
Similar eQTL yield for unannotated expression features compared to annotated features. **a)** Overview of transcriptome quantification. RNA was quantified using five pipelines, each targeting distinct stages of RNA processing, and each followed by eQTL generation. Within annotated regions of the transcriptome, reads were mapped to expression features and thereafter RNA was quantified. These features included: all intronic and exonic regions of a gene (producing gene-intronic gi-eQTLs and gene-exonic ge-eQTLs, respectively); individual exons (producing e-eQTLs) and exon-exon junctions (producing ex-ex-eQTLs). As total RNA was used for library construction, reads mapping to introns were presumed to be due to pre-mRNA within samples (an assumption supported by previous analyses using a subset of these data^41^). Quantification of individual exons and exon-exon junctions provided a means of identifying loci that impact on alternative splicing. In common with most eQTL analyses, we also calculated overall gene expression using all reads mapping to exons of a given gene, resulting in an expression metric which is influenced by transcriptional rate, splicing and RNA degradation rates. Finally, we included annotation-independent approaches to quantify transcription. We focused specifically on unannotated transcription within intergenic regions (producing i-eQTLs, **Online Methods**). **b)** eQTL yields for both tissues were calculated as the number of expression features within a category with at least one significantly associated eQTL divided by the total number of tested features within the same category.

Following stepwise conditional analyses under a false discovery rate (FDR) of 5%, we identified 19,156 separate significant eQTL signals (hereafter “eQTLs”) genome-wide (**Supplementary Tables 2-6**), of which 359 were secondary eQTLs (i.e. eQTLs with independent affects after conditioning on the primary eQTL in the region). While there was a substantial difference in the number of eQTLs identified in putamen and substantia nigra, the difference in sample size (N=111 for putamen and N=69 for substantia nigra) most likely accounted for this. However, eQTL discovery was not simply driven by the number of features tested. Notably, the rate of eQTL discovery, defined as the percentage of all expression features tested with at least one significant associated eQTL, was highest in unannotated intronic and intergenic regions (**Figure 1b**), suggesting that such eQTLs could be biologically important.

### eQTL signals and i-eQTL target regions show high replication rates in independent data sets

We found that 50.6% and 50.4% of the testable eQTLs identified in putamen and substantia nigra respectively were detected using microarray data, generated by the UK Brain Expression Consortium^21^ and based on a common set of RNA samples. We also found that 39.3% of putamen and 50.6% of substantia nigra eQTLs replicated in the GTEx (v7) data resource^22^, using their 111 putamen and 80 substantia nigra samples respectively. Furthermore, we investigated eQTL replication across all brain regions studied in GTEx (**Supplementary Table 7**). We demonstrated a replication rate of 19% to 53.3% for putamen, with the highest replication rates observed in cerebellum (53.3%) and caudate (48.2%). In the case of substantia nigra samples, replication rates were more similar across all brain tissues (50.6% - 62.0%), potentially reflecting the relatively low sample numbers used in the eQTL analysis in both our study (N=65) and that performed by GTEx (N=80). In contrast, when we checked our eQTL signals against those reported by Lappanainen and colleagues^14^ using RNAseq analysis of 373 lymphoblastoid cell lines, we found that despite the larger sample size in this study only 22.0% of putamen and 24.2% of substantia nigra eQTLs were replicated.

Given the paucity of existing eQTL analyses using annotation-independent approaches, we focused on validating the expression of unannotated intergenic regions that were the target of a significant i-eQTL. Using data provided by GTEx and processed for re-use by recount2^23^, we found that 70.3% of all such i-eQTL target regions (in putamen and substantia nigra combined) were detected in at least one other human tissue within the GTEx data set, with the highest validation rates observed amongst brain regions (**Supplementary Table 7**). We also explored the possibility that the transcribed regions detected in our analysis and regulated by i-eQTLs may represent enhancer RNAs as another means of understanding the biological relevance of our findings. Considering all transcribed regions targeted by an i-eQTL and accounting for their genomic size, we found that that there was a 3.0 fold increase in overlap with enhancer regions as defined within the GeneHancer database v4.4^24^ amongst i-eQTL target regions (17.9%) as compared to e-eQTL targets (5.9%) suggesting that the transcribed regions targeted by i-eQTLs are highly enriched for eRNAs.

We further characterised i-eQTL target regions based on their relationship to known genes (**Figure 2a**). Using reads spanning known exons and novel regions, physical proximity and correlation in expression, we categorised unannotated expressed regions into those with strong, moderate or weak evidence for being part of a known gene (**Figure 2a, Online Methods**). This approach allowed us to characterise 68.1% of all unannotated expression regions (**Figure 2b**). The validation rate for expression in the GTEx data resource was 98.5% for unannotated expressed regions with strong evidence for being part of a known gene, 93.8% for those with moderate evidence, and still high at 60.4% for regions with weak evidence for being part of a known gene (**Figure 2c**). We also selected eight unannotated expressed regions for experimental validation (**Supplementary Table 8**). Regardless of their putative relationship to existing genes, all eight regions validated using Sanger sequencing (**Figure 2d**). In the case of unannotated expressed regions with moderate evidence for association, this analysis also enabled us to clarify the exon boundaries. For example, sequencing confirmed the existence of a novel exon of *FIGNL1* (DER18381, **Figure 2d**). Thus, using a combination of public data resources and experimental work, we demonstrated the validity of annotation-independent approaches in transcriptomic analysis.

**Figure 2.**
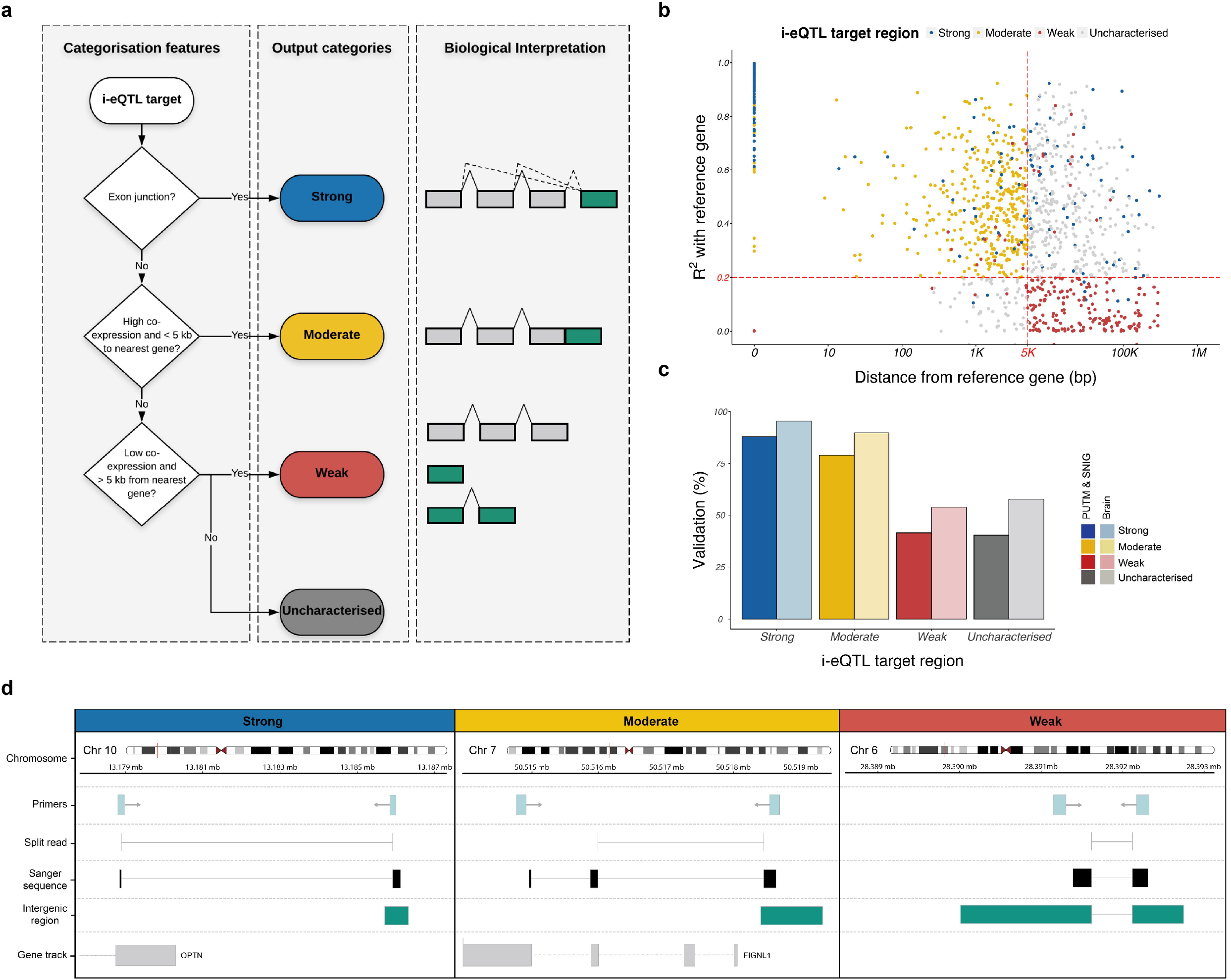
i-eQTL target regions show high replication rates in independent data sets and validate experimentally. **a)** Characterisation of i-eQTL target regions (unannotated expressed regions that were the target of a significant i-eQTL) was based on several features reflecting their relationship to known genes. These features were used to classify these regions into those with strong, moderate and weak evidence for being part of a known gene. Regions categorised as strong and moderate are considered likely to be novel exons of known genes or misannotations of existing exon boundaries, while weak regions are presumed to be independent of any known genes, **b)** Scatterplot of genomic distance and correlation of expression between i-eQTL target regions and their reference genes. **c)** The expression of unannotated expressed regions was validated in GTEx data, using brain region-specific and global brain expression data. Validation rates in putamen and substantia nigra GTEx expression data were combined and displayed separately from validation rates in RNA-seq data from all GTEx brain regions. **d)** Sequencing results for i-eQTL target regions with strong, moderate and weak evidence of being part of a gene. In each case, tracks are provided relating to the location of the primers used to amplify the unannotated expressed region, the RNA-seq split read, the alignment of Sanger-sequenced cDNA, and the predicted boundaries of the unannotated expression region.

### Most i-eQTLs and other non-standard eQTLs represent novel signals

Since 51.2% of our characterised i-eQTL target regions have strong or moderate evidence linking them to a known gene, we further classified i-eQTLs into those with evidence for being new regulatory variants versus those appearing to act in a consistent manner across all exons (thus recapitulating gene-level signals). Using a modified test of heterogeneity^25^ (**Online Methods**) we separately analysed i-eQTLs with strong, moderate and weak evidence for being linked to a known gene (**Figure 3**). This analysis demonstrated that some i-eQTLs were indeed “re-discovered” versions of existing eQTL signals (**Figure 3a**). However, many i-eQTLs appear to be independent regulatory sites. For example, SNP rs4696709 regulates DER10633 expression, a probable novel exon of *ABLIM2* (based on the presence of junction reads), but there is no significant co-regulation of other exons of *ABLIM2* (**Figure 3b**). Even amongst those i-eQTLs with strong evidence linking them to a known gene, the percentage of i-eQTLs sharing signals with known annotation expression features was only 44% (**Figure 3c**). Thus, across all types of characterised i-eQTLs we found evidence for the majority representing novel regulatory variants, acting in a transcript-specific manner.

**Figure 3.**
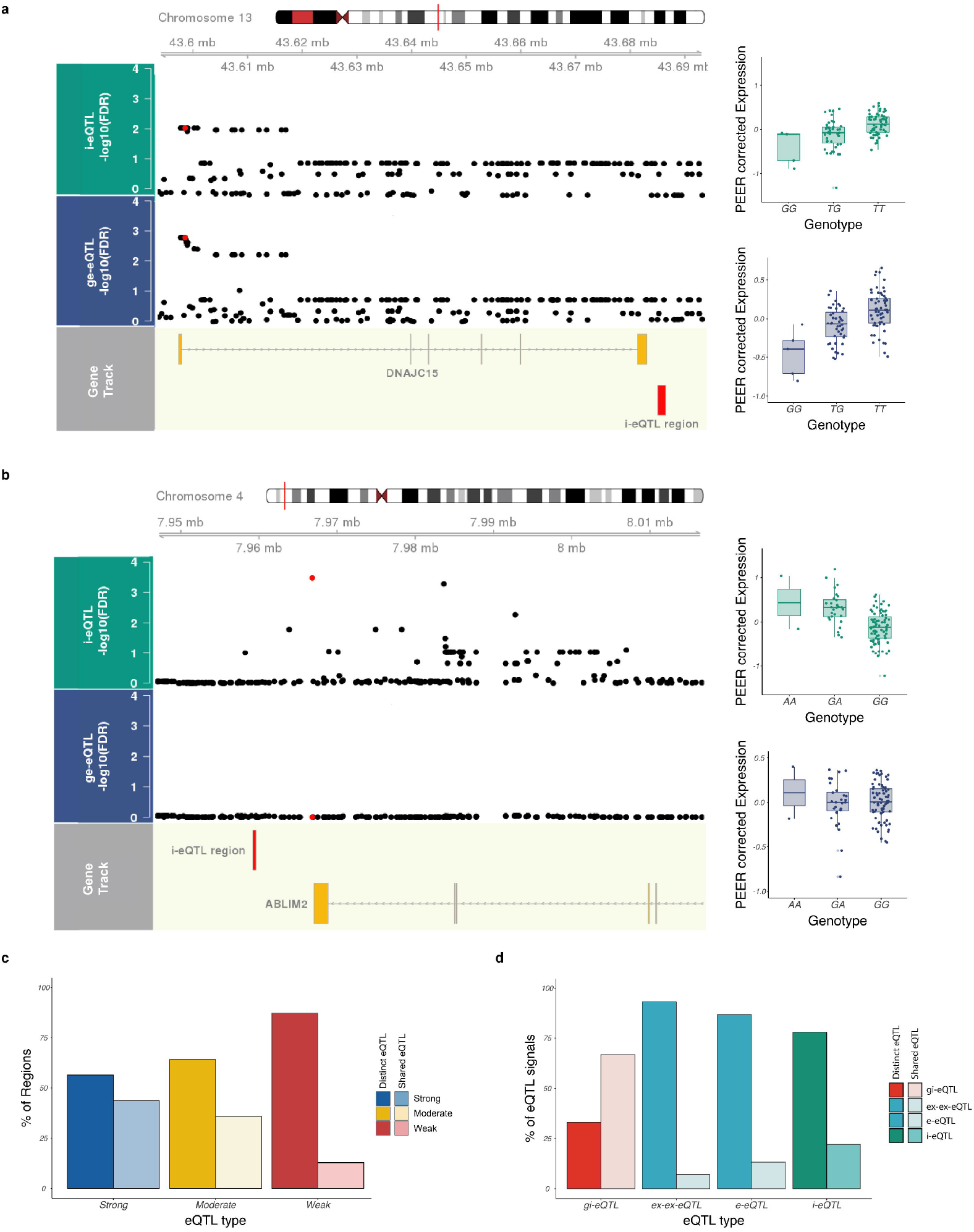
i-eQTL target regions have evidence for distinct regulation. **a)** Local association plots (−log10 FDR-corrected p-values for eQTL association) illustrating sharing of the rs113317084 variant (red point) between the i-eQTL-targeted region, DER32583 (green track), and the ge-eQTL-targeted gene, *DNAJC15* (blue track). **b)** Local association plot illustrating no sharing of the rs4696709 variant (red point) between the i-eQTL-targeted region, DER10633 (green track), and the ge-eQTL-targeted gene, *ABLIM2* (blue track). The detection of reads spanning DER10633 and an annotated exon within *ABLIM2* provides compelling evidence that this region represents a novel exon of the gene. **c)** Heterogeneity (distinct vs. shared) of i-eQTL signals, cross-categorised by the strength of evidence linking their target region to a known gene, suggests that most are distinct and likely represent novel regulatory variants acting in a transcript-specific manner. Heterogeneity was determined using a modified beta-heterogeneity test, accounting for the dependency structure arising from within-individual and within-gene correlations. i-eQTL beta-coefficients were compared to that of the known exon with most evidence of association with the i-eQTL target region. All eQTL signals with an FDR-corrected p-value for heterogeneity < 0.05 were considered distinct, while those with an FDR-corrected p-value > 0.05 were considered shared (similar beta coefficients), **d)** Heterogeneity (distinct vs. shared) of non-standard eQTL classes (gi-eQTLs, e-eQTLs, ex-ex-eQTLs, and i-eQTLs) suggests that many of these classes are distinctly regulated. Heterogeneity was determined using a modified beta-heterogeneity test comparing beta-coefficients from ge-eQTLs to those derived from non-standard eQTL analyses applied to the same gene. This analysis was performed separately for gi-eQTLs (tagging pre-mRNA), e-eQTLs and ex-ex-eQTLs (tagging splicing) and all i-eQTLs (tagging unannotated expression). All eQTL signals with an FDR-corrected p-value < 0.05 were considered distinct, while an FDR-corrected p-value > 0.05 was taken as evidence of eQTL sharing.

We also asked whether our alternative annotation-based eQTL classes (gi-eQTLs, e-eQTLs and ex-ex-eQTLs) provided novel regulatory information compared to the standard gene-level eQTL analysis (ge-eQTLs). Again we used a modified test of beta heterogeneity to determine eQTL signal sharing among these classes. While 66.8% of gi-eQTLs were detectable with “standard” ge-eQTLs, this figure was only 6.9% for splicing eQTLs (**Figure 3d**, suggesting that our additional eQTL classes provided distinct regulatory information driven by splicing effects.

### Splicing eQTLs are enriched for neuronal information

We asked whether our different eQTL classes varied in terms of the cellular specificity of their target expression features. We used weighted gene co-expression network analysis, in combination with publicly available cell-specific annotation data, to assign eQTL target expression features to one of five broad cell types: neuron, oligodendrocyte, astrocyte, microglia and endothelial cell (**Figure 4a, Online Methods**). This module-membership approach allowed us to provide putative cellular classifications for expression features even if they were outside known annotations. We confidently assigned up to 75% of all analysed genes to a specific cell type, and these were then related to 41.5% of all eQTL target expression features. We observed a significant enrichment of neuronal genes in all non-standard eQTL classes in one or both tissues investigated (**Figure 4b, Supplementary Table 8**). These included i-eQTLs targeting unannotated expressed regions (FDR-corrected p-value = 1.20 × 10^−2^ in putamen, **Figure 4b**). Furthermore, we found that the targets of splicing eQTLs were significantly enriched for neuronal genes (FDR-corrected p-values = 1.21 × 10^−7^ and 2.28 × 10^−5^ for e-eQTLs and ex-ex-eQTLs respectively in substantia nigra), oligodendrocyte genes (FDR-corrected p-value = 1.7 × 10^−3^ and 4.1 × 10^−2^ for e-eQTLs and ex-ex-eQTLs respectively in substantia nigra) and astrocyte genes (FDR-corrected p-value = 8.74 × 10^−4^ and 1.12 × 10^−3^ for e-eQTLs and ex-ex-eQTLs respectively in substantia nigra, **Figure 4b**). This points to the importance of capturing splicing information in the analysis of human brain samples.

**Figure 4.**
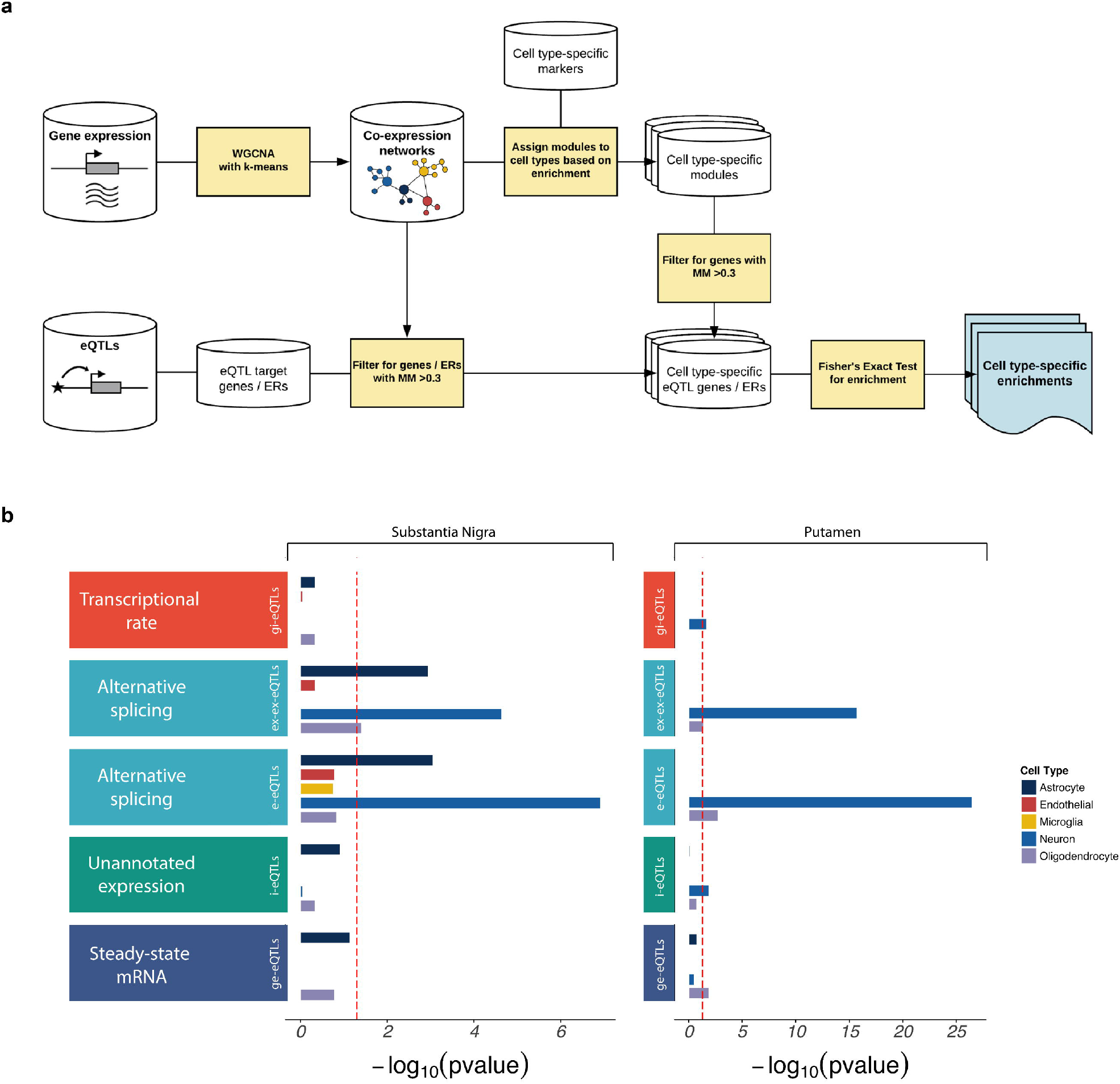
Non-standard eQTL analyses produce additional biologically-relevant information. **a)** Schematic diagram showing the use of gene co-expression networks to assign eQTL target genes and unannotated expressed regions (ERs) to the cell type most likely to be driving gene expression in the tissue. We used the WGCNA R package^42^ refined with a k-means algorithm^43^ to construct separate gene co-expression networks for each tissue separately. We annotated modules for cell type-specific enrichments using cell type-specific marker genes sets^44–47^,⍰. Genes assigned to modules significantly enriched for brain-related cell type markers and with a module membership of > 0.3 were allocated a cell type “label”. Next, for each eQTL targeting a known genic region or an unannotated expressed region with high or moderate evidence linking it to a known gene, if the target gene was allocated to a cell type then the related eQTL received the same cell type label. For eQTLs targeting unannotated expressed regions with low evidence for association with a known gene or which could not be classified, we assigned the target expression feature to a module (and by inference a cell type) based on its highest module membership providing the module membership was at least 0.3. Finally, for each eQTL class and each cell type, namely neuron, microglia, astrocyte, oligodendrocyte and endothelial cell, we applied a Fisher’s Exact test to test for enrichment of that cell type label amongst the genes associated to the eQTL class. **b)** Expression features targeted by different eQTL classes were variably enriched for genes with cell-biased expression, highlighting the importance of capturing this information. Enrichment of genes with cell-biased expression within eQTL targeted expression features was performed separately for each tissue and was determined using a Fisher’s Exact test and a significance cut-off of P < 0.05 (dashed red line at −log_10_(P) = 1.30). Genes assigned to modules significantly enriched for brain-related cell type markers and with a module membership of > 0.3 were allocated a cell type “label”. Next, for each eQTL targeting a known genic region or an unannotated expressed region with high or moderate evidence linking it to a known gene, if the target gene was allocated to a cell type then the related eQTL received the same cell type label. For eQTLs targeting unannotated expressed regions with low evidence for association with a known gene or which could not be classified, we assigned the target expression feature to a module (and by inference a cell type) based on its highest module membership providing the module membership was at least 0.3. Finally, for each eQTL class and each cell type, namely neuron, microglia, astrocyte, oligodendrocyte and endothelial cell, we applied a Fisher’s Exact test to test for enrichment of that cell type label amongst the genes associated to the eQTL class. **b)** Expression features targeted by different eQTL classes were variably enriched for genes with cell-biased expression, highlighting the importance of capturing this information. Enrichment of genes with cell-biased expression within eQTL targeted expression features was performed separately for each tissue and was determined using a Fisher’s Exact test and a significance cut-off of P < 0.05 (dashed red line at −log_10_(P) = 1.30).

### eQTLs targeting unannotated transcribed regions are enriched for disease-relevant information

We investigated the overlap of unannotated transcribed regions and eQTL sites with known GWAS association signals. We used the US National Human Genome Research Institute (NHGRI) GWAS catalogue, restricted to genome-wide significant SNPs (*P* < 5 × 10^−8^) and stratifying for SNP-phenotype associations of relevance to neurological/behavioural disorders as defined within the STOPGAP database^26^. First, we investigated the possibility that unannotated transcribed regions could themselves harbour risk loci of relevance to neurological diseases, by calculating the proportion of transcribed intergenic regions containing a brain-relevant risk locus and comparing this value to that for expressed exons. After adjusting for the size of each annotation, we found that unannotated transcribed regions and exons had a similar level of overlap with brain-relevant risk loci (3.5% for exons and 2.9% for transcribed intergenic regions after adjustment for annotation size). However, the enrichment of brain-relevant risk loci was higher for novel transcribed regions as compared to exons (1.51 fold for targets of e-eQTLs versus 2.70 fold for targets of i-eQTLs after adjustment for annotation size). Furthermore, we found a significant enrichment for GWAS variants that were associated with neurological and behavioural disorders compared to all other SNP-phenotype associations (**Supplementary Figure 2**) amongst our eQTLs. While the enrichment of brain-relevant GWAS associations was most evident in ex-ex-eQTLs and gi-eQTLs, we also found a significant enrichment for i-eQTLs (FDR-corrected Fisher’s Exact p-value = 6.45 × 10^−7^). i-eQTLs provided useful information for 36.7% of all the neurologically-relevant risk loci within this analysis (equating to 76 loci). Given that these findings could potentially be driven by correlations between i-eQTLs and more conventional eQTL signals, we repeated this analysis only using i-eQTLs considered to be independent regulatory signals (based on modified beta-heterogeneity testing described above). We found that the enrichment of brain-relevant risk loci amongst i-eQTLs increased in significance in this sub-group (FDR-corrected p-value= 7.01 × 10^−8^). Thus, our analysis suggests that i-eQTLs do contribute to the understanding of a significant proportion of neurologically-relevant risk loci.

We further explored signal enrichment in i-eQTLs in relation to two neurological diseases related to basal ganglia dysfunction: Parkinson’s disease and schizophrenia. Using GWAS summary statistics for these diseases^1,11^, we performed colocalisation analyses for disease-risk association signals against i-eQTL signals using the coloc^26^ program. We identified twenty-three i-eQTL signals that colocalised with risk loci for schizophrenia or Parkinson’s disease (**Supplementary Table 10**). Amongst the former, we identified a signal indexed by the lead SNP rs35774874 (GWAS p-value = 1.97 × 10^−11^) that colocalised with an i-eQTL targeting a probable novel 3’UTR of *SNX19* (posterior colocalisation probability = 0.75), a gene which has already been highlighted in schizophrenia^27,28^. Similarly we identified a colocalisation of the PD GWAS lead SNP rs4566208 (GWAS p-value = 2.28 × 10^−7^) with an i-eQTL regulating a probable novel exon of *ZSWIM7* (i-eQTL p-value =1.09 × 10^−5^; posterior colocalisation probability = 0.89, **Supplementary Figure 3**). However, we also found seven co-localising i-eQTL signals targeting unannotated expressed regions that were not linked to a known gene. For example, we found that the schizophrenia risk SNP rs12908161 (GWAS p-value = 9.41 × 10^−10^) had a posterior colocalisation probability of 1.00 with an i-eQTL targeting the unannotated expressed region DER36302 (chr15:84833811-84833975, eQTL p-value = 7.94 × 10^−10^ in putamen, **Figure 5a**). This novel transcribed region appears to be independent of neighbouring genes, and is expressed in human brain, with the highest expression in the anterior cingulate and frontal cortex, two brain regions relevant to schizophrenia (**Figure 5b**).

**Figure 5.**
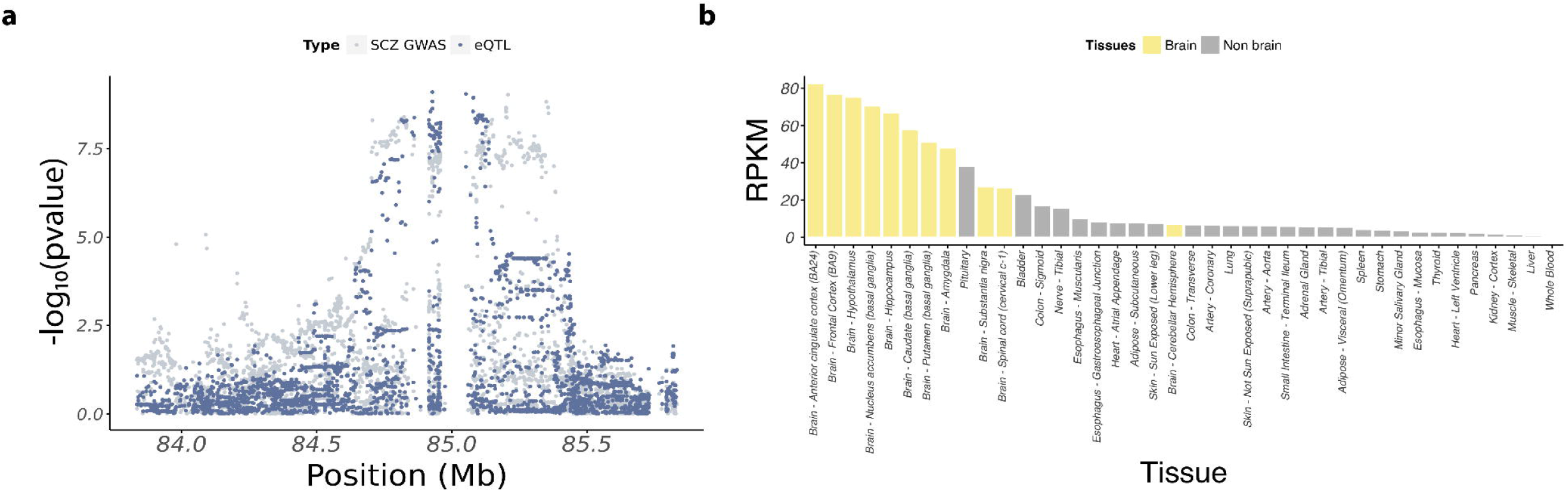
Annotation-independent approaches produce disease-relevant information unavailable through annotation-dependent approaches. **a)** Colocalisation of the schizophrenia GWAS lead SNP rs950169 (GWAS p-value = 7.62 × 10^−11^) and the i-eQTL targeting DER36302 (eQTL p-value = 1.15 × 10^−10^ in putamen). **b)** Expression of DER36302 across tissues sampled by the GTEx consortium. Brain tissues are highlighted in yellow with the anterior cingulate and frontal cortex showing the highest expression.

### ASE discovery and validation

We applied allele-specific expression (ASE) analysis to a subset of 84 brain samples (substantia nigra n=35; putamen n=49) for which we had access to whole exome sequencing in addition to SNP genotyping data (**Online Methods, Supplementary Table 1**). ASE analysis quantifies the variation in expression between two haplotypes of a diploid individual distinguished by heterozygous genetic variation, and so can capture the effects of a range of regulatory processes, namely genomic imprinting, nonsense-mediated decay and cis-regulation (**Figure 6a**). In total 252,742 valid heterozygous SNPs (hetSNPs) across 53 individuals were analysed. Of these, 7.62% (19,266) were significant ASE signals (hereafter “ASEs”) at FDR <5% in at least one sample, covering 8,654 genes. Of the 19,266 ASEs identified, 12,096 were found in putamen and 11,871 in substantia nigra (**Supplementary Table 11**). Consistent with previous studies, we found that ASEs can operate as markers of imprinting or parent-of-origin effects: ASE signals that are not unidirectional across individuals are expected to be enriched for imprinted genes (**Figure 6a**). Consistent with expectation, of all genes containing an ASE, 170 were identified on the X chromosome. Furthermore, we observed that all inconsistent ASE signals (those that were not unidirectional within >= 10 individuals) were located within genes known to be imprinted (as reported in www.geneimprint.com or within the literature^27–30^) compared to 1-5% of consistent signals (**Figure 6b**).

**Figure 6.**
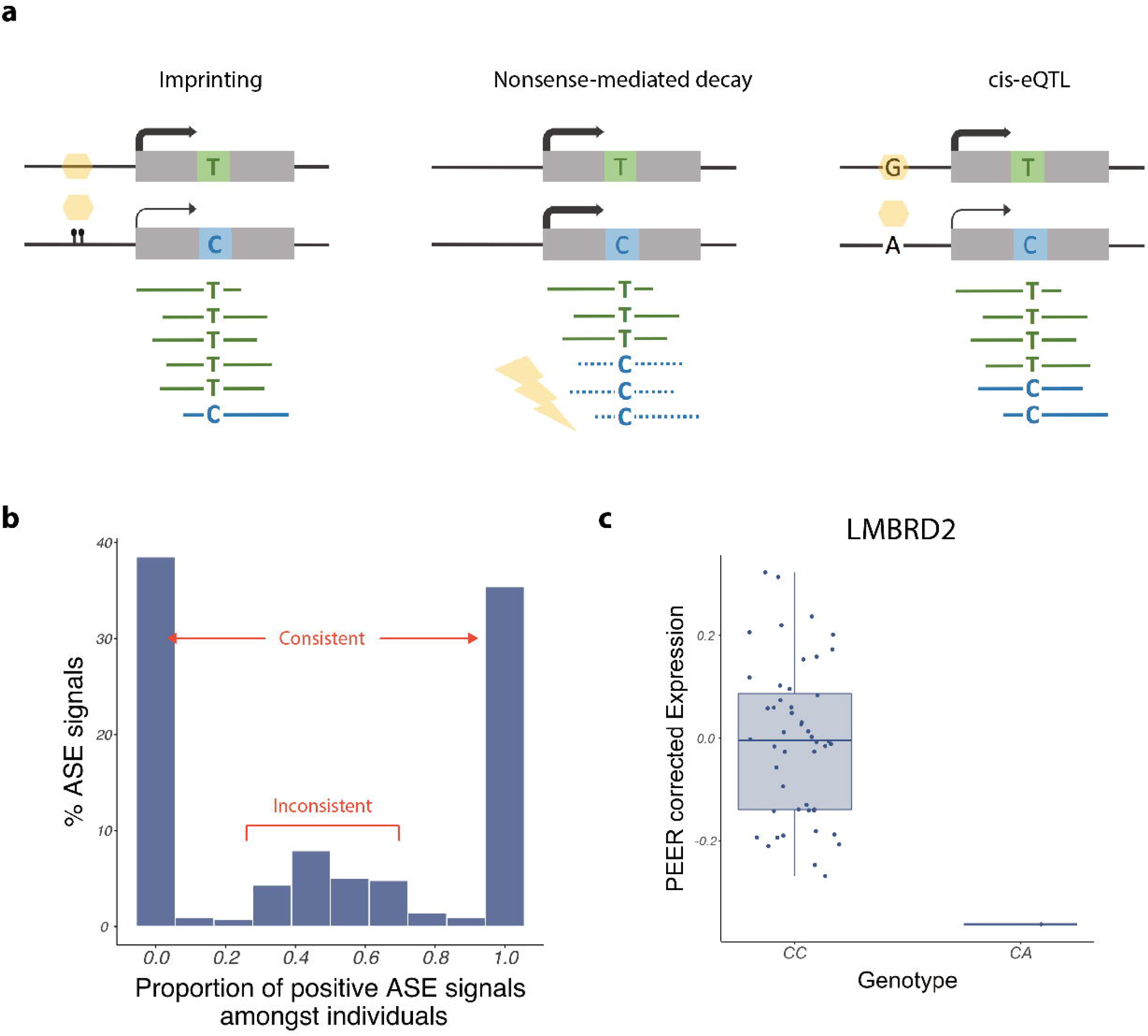
Allele-specific expression provides evidence of dosage compensation in human brain-expressed genes. **a)** Overview of the mechanisms by which allele-specific expression can arise. Allele-specific expression can arise through epigenetic effects (e.g. imprinting), heterozygous mutations triggering nonsense-mediated decay of transcripts, and regulation by (for example) a cis-regulatory variant (cis-eQTL). **b)** The majority of allele-specific expression signals passing FDR < 0.05 produced unidirectional signals (0 or 1) and were considered consistent. Inconsistent ASE signals (those that were not unidirectional in ≥ 10 individuals) were found only in known imprinted genes, thus providing additional validation of our ASE signals. **c)** Comparison of *LMBRD2* expression in putamen from one individual heterozygous for a rare stop gain mutation (CA) in the gene versus all other individuals (CC) revealed a significant reduction in *LMBRD2* expression, implying effective nonsense-mediated decay. Data presented using Tukey-style box plots.

We also found evidence for the generation of ASE signals through nonsense-mediated decay. We identified 61 protein-truncating variants (defined as stop gain, donor splice site and donor acceptor mutations) among our ASEs. Consistent with expectation, the majority of these variants were predicted to cause nonsense-mediated decay (52.5% using SNPeff^31^) and appeared to result in mono-allelic expression, with >95% of all reads at the ASE site originating from a single allele. These extreme ASEs are expected to generate an effective reduction in gene dosage, and therefore to cause a significant reduction in the total expression of the affected genes. An example of this pattern is seen in the *LMBRD2* gene (**Figure 6c**).

Finally, to check the overall reliability of our findings, we looked for validation of our ASEs in an independent data set of 462 lymphoblastoid cell lines reported by Lappanainen and colleagues^14^. We found that 67% of testable ASE signals could be detected at an FDR <5%, demonstrating the reliability of our ASE sites while also suggesting the presence of brain-specific ASEs.

### ASEs tag both gene level and splicing eQTLs

Cis-regulatory variants are known to be one important generator of ASEs^32^. We therefore investigated the overlap of eQTLs with ASEs in our data, and compared it to the overlap observed with randomly selected non-ASE heterozygous SNPs. After controlling for read depth, we identified a highly significant enrichment of eQTLs amongst our ASEs (p-value = 2.65 × 10^−195^ in putamen and 9.99 × 10^−111^ in substantia nigra). This enrichment remained significant when we restricted our analysis to eQTLs with effects on splicing (e-eQTLs and ex-ex-eQTLs, p-value = 1.17 × 10^−178^ in putamen and 1.29 × 10^−89^ in substantia nigra). However, as expected, it was absent when we considered ASEs located within imprinted genes, where the parental origin of the SNP rather than the impact of cis regulatory sites is expected to drive allele-specific expression (p-value = 0.923 in putamen and 0.856 in substantia nigra).

To investigate the extent to which ASE sites tagged gene-level or transcript-specific cis-regulatory effects, we focussed on common ASE sites (seen in >=10 individuals) that were unidirectional in nature (same direction of effect across all individuals). For each valid ASE site, we measured exon expression across all three genotypes. Of all testable ASEs, we found that 43.2% were also likely to be eQTLs. To ask whether the underlying cis regulatory effects operated in an exon-specific or gene-level manner, we repeated the analysis using gene-level expression across the genotypes. Of the ASEs that were also likely eQTLs, 51.8% appeared to operate in an exon-specific manner, implying that they tagged splicing eQTLs. rs7724759, a splice site variant present in the *CAST* gene (**Figure 7a and Figure 7b**), and rs1050078, a variant in *SNX19* (**Figure 7c and Figure 7d**), are example of ASEs likely to be driven by exon-level and gene-level regulation respectively.

**Figure 7.**
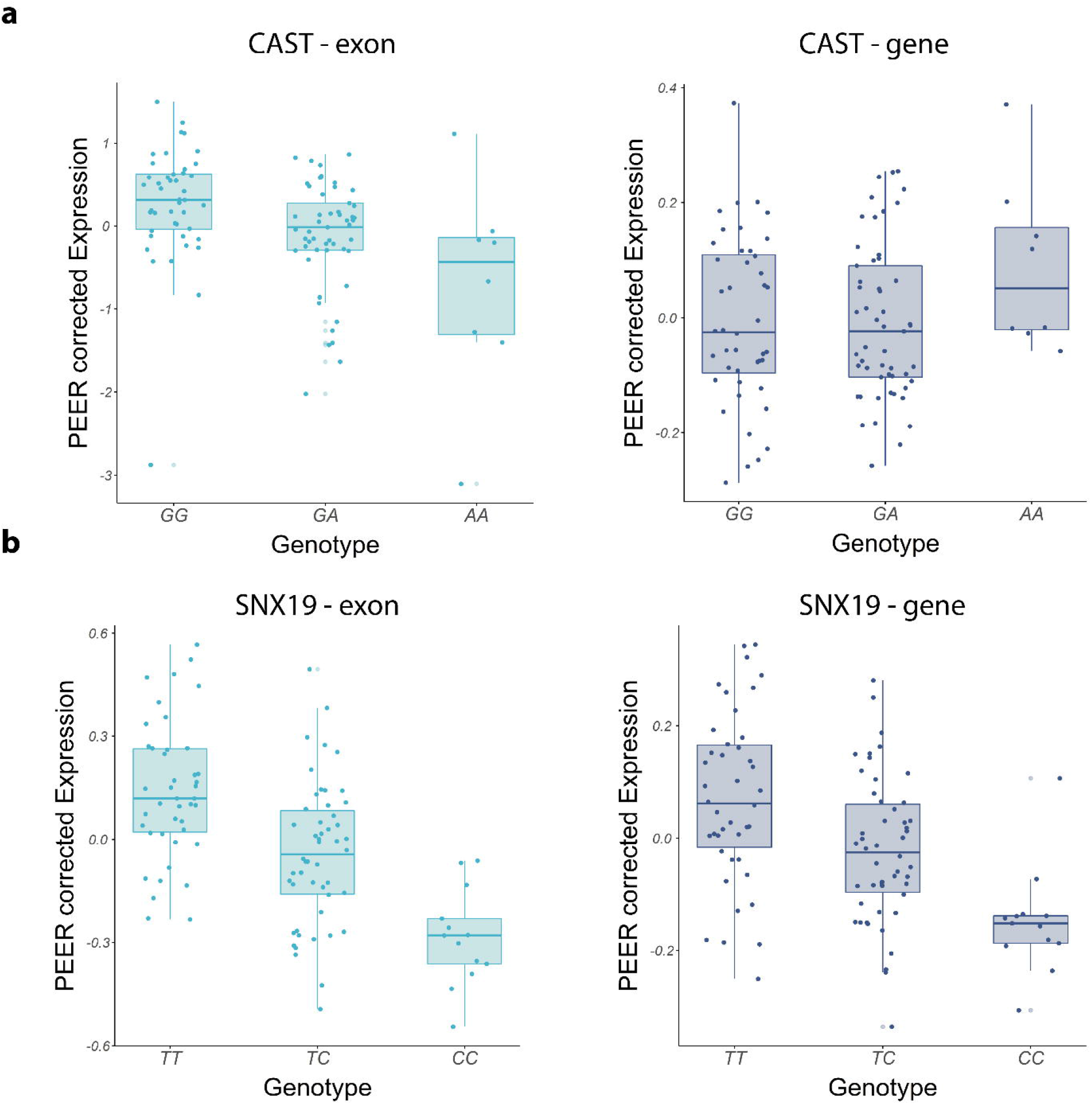
Allele-specific expression sites capture both splicing and gene level cis-regulation. **a)** Some ASEs appear to operate in an exon-specific manner, and likely represent splicing QTLs. Expression of the exon containing rs7724759, a splice variant present in the *CAST* gene, across individuals of all three genotypes demonstrates that the dosage of the splice variant impacts on exon expression. However, *CAST* gene-level expression across individuals of all three genotypes is unaffected by the dosage of the splice variant. **b)** Some ASEs appeared to operate in a gene-level manner. Expression of the exon containing rs1050078, a variant present in the *SNX19* gene, and *SNX19* gene-level expression across individuals of all three genotypes demonstrates a similar dosage relationship. Data presented using Tukey-style box plots.

While this approach allowed us to identify ASE sites tagging splicing eQTLs, it was limited to a small subset of common ASE sites and represented only 0.8% of all ASEs. To address this issue, we used the machine learning program SPIDEX to predict the effect of all ASEs on splicing^33^. Given a genetic variant, SPIDEX provides the delta percent inclusion ratio (ΔΨ) for the exon in which the variant is located (reported as the maximum ΔΨ across tissues). We compared predicted ΔΨ values at ASE sites versus non-ASE sites and found significantly higher values amongst ASEs (Fishers test p-value = 4.50 × 10^−5^ and p-value = 2.07 × 10^−19^ using a randomization approach – see Online Methods). This strongly suggests that ASEs are enriched for variants with effects on splicing.

### ASEs are significantly enriched for biologically- and disease-relevant information

We assessed the cellular specificity of the regulatory information ASEs provide. Given that ASE analysis is performed within an individual and so is not subject to the confounding effects of cellular heterogeneity across individuals, we expected that ASEs would be a powerful means of obtaining cell-specific regulatory information. Using a similar approach to that applied to eQTLs to assign genes containing ASEs to brain-relevant cell types (neurons, oligodendrocytes, astrocytes, microglia and endothelial cells), we found that ASE-containing genes were highly enriched for neuronally–expressed genes (FDR-corrected p-values of 9.97 × 10^−235^ in putamen and 3.05 × 10^−97^ in substantia nigra). We also found significant enrichments (FDR p-value <5%) for oligodendrocyte, astrocyte, microglia and endothelial gene sets. While we observed a similar pattern of cell-type specific enrichments amongst eQTLs, the strength of evidence for cellular specificity of ASEs was striking, suggesting that incomplete covariate correction may be hampering the power of eQTL analyses (**Figure 8a, Supplementary Table 9**).

**Figure 8.**
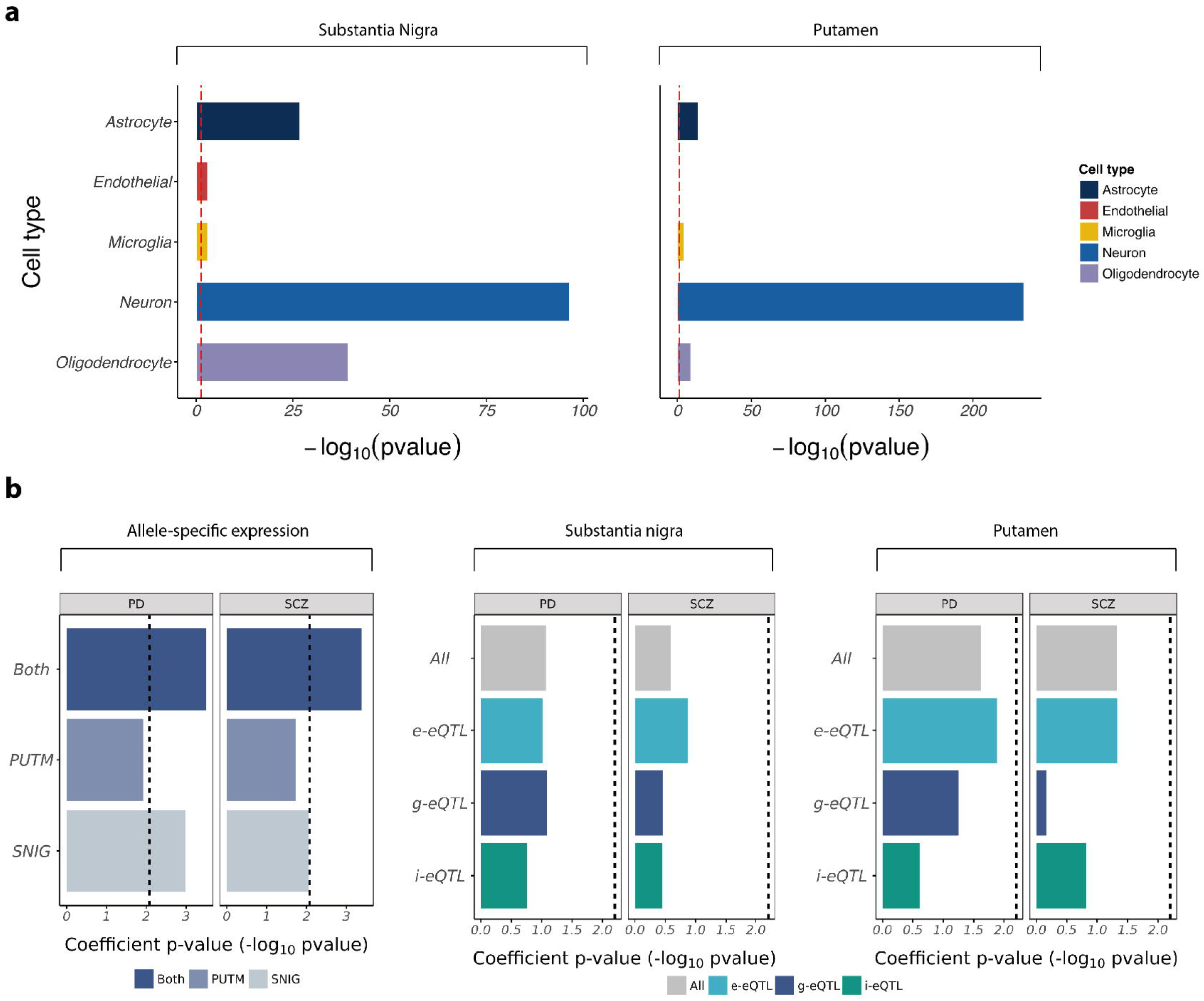
Allele-specific expression sites biologically- and disease-relevant information. **a)** Genic locations of ASEs were highly enriched for genes with cell-biased expression in both putamen and substantia nigra. Enrichment of genes with cell-biased expression within ASE locations was performed separately for each tissue and was determined using a Fisher’s Exact test and a significance cut-off of p-value < 0.05 (dashed red line at −log_10_(P) = 1.30). **b)** Enrichment of heritability for Parkinson’s Disease and schizophrenia in ASEs and eQTLs identified in substantia nigra, putamen and across both tissues.

Finally, we used GWAS summary data sets for Parkinson’s disease^1^ and schizophrenia^11^ to investigate the disease-relevance of ASEs. Since GWAS loci often lie close to genic regions and so are likely to overlap by chance with ASE signals, we used a randomization approach^34^ to investigate the enrichment of GWAS loci within our ASEs (**Online Methods**). We compared overlaps between risk loci and ASEs to overlaps between risk loci and randomly selected non-ASE sites, and found a highly significant enrichment of GWAS risk loci for both schizophrenia (p-value = 7.49 × 10^−35^ for ASEs derived from both tissues) and Parkinson’s disease (p-value = 4.19 × 10^−7^ for ASEs derived from both tissues). We validated these findings using stratified LD score regression^35^ by treating our ASE sites as a form of binary annotation, and interestingly found that the enrichment of PD heritability appeared to be more specific to ASEs identified in substantia nigra using this approach (**Figure 8b, Supplementary Table 12**). There was no enrichment in PD or schizophrenia heritability amongst eQTLs using the same method. Thus, while we recognise that eQTLs can be powerful when linked to even more specific cell types for this type of analysis^36,37^, we demonstrate the additional power of ASE analysis to generate disease-relevant information, despite the small number of samples we had at our disposal.

## Discussion

The human brain is an especially challenging organ in which to conduct eQTL and ASE studies. In addition to the difficulties of sample collection, the brain is a highly complex organ, with site-specific pathologies that motivate the use of equivalently specific analyses. The brain is also known to express many transcripts not seen in other parts of the body, and it is suspected that much of its transcriptome remains uncharacterised^20,38^.

We tackled these issues by collecting RNA-seq data from human substantia nigra and putamen, and applied a bank of five transcriptome quantification methods, including annotation-agnostic approaches as well as approaches to interrogate different stages of RNA processing. We found that there is significant variation between eQTL classes in their neuron- and brain-specific information content, as measured by the cell type-specific enrichment of eQTL targets. The most neuronally-enriched and brain-specific results were found in eQTL classes that most closely tag the regulation of splicing (e-eQTLs and ex-ex-eQTLs) rather than gene level expression. This finding is consistent with recent studies that suggest that splicing eQTLs can provide significant insights into complex diseases in general^39^, and brain-related disorders in particular (e.g. schizophrenia^36,37^). Thus, in addition to providing a rich eQTL resource, our study suggests that the utility of existing and future eQTL analyses in human brain may critically depend on the ability of the RNA sequencing technology, and of the analytic methods applied, to capture transcript-specific information.

We also asked whether the incomplete annotation of the human brain transcriptome might be limiting eQTL discovery as well as reducing the tissue-specific nature of the regulatory information discovered. We focused on transcription within intergenic regions, since transcriptional activity in these regions cannot be explained through the presence of pre-mRNA but could be generated through the expression of long intergenic non-coding RNAs and enhancer RNAs, as these are reported to be expressed in a highly tissue specific manner^19,40^. We show that these expressed intergenic regions are reliably detected, and that approximately 16.1% of these expressed regions are highly likely to represent novel exons of known genes (as demonstrated through the existence of junction spanning reads). They are also enriched for overlap with enhancer regions, suggesting that many could also represent eRNAs (3.03-fold enrichment over expressed exons). Finally, we show that intergenic eQTLs (i-eQTLs) are enriched for neuronally-relevant information, and most importantly that they can provide unique disease insights that would be missed using standard analyses, as illustrated by the colocalisation of i-eQTL signals with schizophrenia risk loci.

Nevertheless, the identification of splicing eQTLs from homogenates of macro-dissected human brain, particularly from brain regions which are hard to obtain in large numbers, is likely to remain challenging even after accounting for the on-going development of tools to optimise transcriptome quantification. This motivates the use of ASE analysis, a form of within individual comparison that compares variation in expression between two haplotypes of a diploid individual. This within-individual comparison means that ASE analysis is unaffected by between-individual confounders, such as the variability in cell type-specific density among individuals. We applied ASE analysis to 49 putamen and 35 substantia nigra samples, for which both whole exome sequencing and genotyping data were available. Consistent with our expectation, we found that ASEs were significantly enriched for variants identified as splicing eQTLs within our own analysis or as predicted to affect splicing according to SPIDEX. Furthermore, we found that the ASEs we identified tagged regulatory information that was highly enriched for neurons and brain-relevant cell types, even after accounting for the general enrichment in brain-specific information contained within the RNA-seq data. Finally, and most importantly, we used two separate approaches to demonstrate the relevance of ASEs to complex forms of both Parkinson’s disease and schizophrenia, with evidence for enriched heritability amongst ASEs. Given the small numbers of samples used in our ASE analyses, this finding is particularly important. Thus, we provide evidence to suggest that ASE analysis may be a particularly effective and efficient means of obtaining regulatory information relevant to splicing, cell-type and disease.

In summary, by using a range of methods to quantify and analyse brain transcriptomic data, we demonstrate the importance of capturing information on the regulation of known and novel splicing for the understanding of complex brain disorders, and show that more effective ASE analyses performed even on small sample sets can provide additional insights.

## Supporting information

Suppmentary Figure 1

Suppmentary Figure 2

Suppmentary Figure 3

Suppmentary Figure 4

## Supplementary Figures

Supplementary Figure 1

Screenshots to show the information accessible through the BRAINEAC web resource a) Use of BRAINEAC to access eQTL data. b) Use of BRAINEAC to access gene co-expression network data.

Supplementary Figure 2

Enrichment of risk loci for neurological and behavioural disorders across all eQTL classes.

Supplementary Figure 3

Colocalisation of the PD GWAS lead SNP SNP rs4566208 (GWAS p-value = 2.28 × 10^−7^) and the i-eQTL targeting DER38036 (i-eQTL p-value =1.09 × 10^−5^ in putamen).

Supplementary Figure 4

Heatmap to show the relationship between PEER axes (X-axis) and known covariates (Y-axis). The FDR-corrected p-values for correlations between each PEER axis and known factor are depicted by the colour of ea. The Pearson R^2^ values are displayed within each cell.

## Supplementary Tables

Supplementary Table 1

Table summarising brain samples used for the generation of RNAseq data

Supplementary Table 2

Table of gene-intronic eQTLs (gi-eQTLs) at FDR <5%

Supplementary Table 3

Table of exonic eQTLs (e-eQTLs) at FDR <5%

Supplementary Table 4

Table of exon-exon junction eQTLs (ex-ex-eQTLs) at FDR <5%

Supplementary Table 5

Table of gene-exonic eQTLs (ge-eQTLs) at FDR <5%

Supplementary Table 6

Table of intergenic eQTLs (i-eQTLs) at FDR <5%

Supplementary Table 7

Table of eQTL replication rates across all GTEx brain-specific eQTL datasets

Supplementary Table 8

Table of i-eQTL target region validation

Supplementary Table 9

Cell type-specific enrichments of eQTL target regions

Supplementary Table 10

i-eQTLs colocalising with GWAS loci for schizophrenia and Parkinson’s Disease

Supplementary Table 11

Table of ASEs identified in putamen and substantia nigra with a MAF > 5%

Supplementary Table 12

Enrichment of PD and schizophrenia heritability amongst ASEs using stratified LD score regression

## Acknowledgements

Mina Ryten, David Zhang and Karishma D’Sa were supported by the UK Medical Research Council (MRC) through the award of Tenure-track Clinician Scientist Fellowship to Mina Ryten (MR/N008324/1). Sebastian Guelfi was supported by Alzheimer’s Research UK through the award of a PhD Fellowship (ARUK-PhD2014-16). Regina Reynolds was supported through the award of a Leonard Wolfson Doctoral Training Fellowship in Neurodegeneration.

All RNA sequencing data performed as part of this study was generated by the commercial company AROS Applied Biotechnology A/S (Denmark).

## Author contributions

Sebastian Guelfi processed the RNA sequencing data and generated the eQTL data. Karishma D’Sa generated the ASE data and with Juan Botia developed and implemented the randomization analysis. Juan Botia also performed all network-based analyses. Jana Vandracova assisted with the annotation of eQTLs and ASEs. David Zhang performed coloc analyses. Regina Reynolds validated novel transcribed regions and with the assistance of Sarah Gagliano conducted analyses using LD score regression. Adaikalavan Ramasamy performed the analysis of the genotyping data and contributed to the development of the ASE and eQTL pipelines. Daniah Trabzuni performed the RNA extractions from post-mortem human brain tissue. Leonardo Collado-Torres assisted with the analysis of data available through recount2. Andy Thomason and Pedro Quijada Leyton designed and generated the web server to enable access to eQTL and ASE data generated within this study. Mike Nalls provided assistance with the use of the Parkinson’s Disease GWAS data. Kerrin Small provided input regarding the development of the randomization analysis. Colin Smith assisted with the collection of human putamen and substantia nigra samples together with the UK Brain Expression Consortium. Michael Weale and Mina Ryten supervised the generation of the ASE and eQTL pipelines. Sebastian Guelfi, Karishma D’Sa, Juan Botia, Jana Vandracova, Michael Weale, John Hardy and Mina Ryten prepared the manuscript. Michael Weale, John Hardy and Mina Ryten conceived and designed the project.

