## Supplementary material for "Regulatory sites for known and novel splicing in human basal ganglia are enriched for disease-relevant information": Online Methods

### Generation and processing of RNA sequencing data

Human brain samples originating from 117 individuals of European descent were obtained from the Medical Research Council (MRC) Sudden Death Brain and Tissue Bank and the Sun Health Research Institute. All samples were authorised for ethically approved scientific investigation (Research Ethics Committee number 10/H0716/3) and had fully informed consent for retrieval. These samples constituted a subset of the United Kingdom Brain Expression Consortium (UKBEC) data set and a detailed description of tissue processing, dissection and RNA extraction has been reported previously^1^^,^^2^. In brief, the miRNeasy 96 sample kit (Qiagen, UK) was used to isolate total RNA, and the RNA integrity number was assessed for each sample using the Agilent 2100 Bioanalyzer (Agilent Technologies UK Ltd, UK) in combination with the RNA 6000 Nano-LabChip kit (**Supplementary Table 1**).

cDNA libraries were prepared by the UK Brain Expression Consortium in conjunction with AROS Applied Biotechnology A/S (Aarhus, Denmark) as previously reported^3^. In brief, 100 ng total RNA was used as input for cDNA generation using NuGen’s Ovation RNA-seq System V2 (NuGen Technologies, US). The RNA was processed according to the manufacturer’s protocol resulting in amplified cDNA from total RNA and concomitant de-selection of rRNA. Importantly, reverse transcription in this protocol is carried out using both oligo dT and random primers. This allowed total RNA profile patterns to be assessed with the latter. 1μg of the cDNA was fragmented using a Covaris S220 Ultrasonicator and the fragmented cDNA was used as the starting point for Illumina’s TruSeq DNA library preparation (Illumina, US). Finally, library molecules containing adapter molecules on both ends were amplified through 10 cycles of PCR. The libraries were sequenced using Illumina’s TruSeq V3 chemistry / HiSeq2000 and 100 base pair paired-end reads. The sequencing data was converted to fastq-files using Illumina’s CASAVA Software. Paired end data was mapped to the human genome (build GRCh37) using tophat2^4^ (v2.0.9) with default settings and a transcriptome-guided approach using the Ensembl reference (v72) based on GENCODE version 18. Reads mapping to rRNA regions were removed from the analysis.

Transcriptome quantification was performed using the transcriptome definition from Ensembl reference (v72) using HTSeq-Counts^5^ (v0.5.4p4) for exonic regions, BEDtools^6^ for intronic regions, DEXSeq^7^ (v1.10.6) for individual exons, Altrans^8^ (v1.1.02) for exon-exon junctions and the derfinder R package^9^ for unannotated transcribed regions. Expressed regions were identified by derfinder using default settings and filtering for expressed regions of >100bp in length, annotated as intergenic based on Ensembl v72 and UCSC through the R library TxDb.Hsapiens.UCSC.hg19.knownGene version v3.1.2, and which uniquely mapped (defined as > 98% alignment precision). This approach was applied to each tissue separately (putamen and substantia nigra). All forms of transcriptome quantification were normalised using CQN^10^, with GC content calculated separately for each type of quantification and used as an input for the CQN R-bioconductor package. The resulting normalised expression data was then transformed into RPKM values and log2 converted. Exonic, intronic, and transcribed intergenic regions with an RPKM of greater 0.1 in at least 80% of samples were selected for downstream analyses.

### DNA extraction and genotyping

As previously reported, genomic DNA was extracted from sub-dissected samples (100–200 mg) of human postmortem brain tissue using Qiagen’s DNeasy Blood & Tissue Kit (Qiagen, UK). All samples were genotyped on the Illumina Infinium Omni1-Quad BeadChip and on the Immunochip, a custom genotyping array designed for the fine mapping of autoimmune disorders. After standard quality controls as previously reported, imputation was performed using MaCH^11^ and minimac^12^ using the European panel of the 1000 Genomes Project (March 2012: Integrated Phase I haplotype release version 3, based on the 2010 November data freeze and 14 March 2012 haplotypes). We used the resulting 5,878,211 SNPs and 576,942 indels with good postimputation quality (R^2^ > 0.50) and minor allele frequency (MAF) of at least 5%.

A subset of DNA samples were also analysed using whole exome sequencing as previously reported^13^ (N=57, **Supplementary Table 1**, EGAS00001002113). Exome sequencing was performed on each subject of the cohort according to the manufacture’s capture protocol; captures included Illumina and Nimblegen. Paired end sequence reads were aligned with BWA against the reference human genome (UCSC hg19)^14^. Duplicate read removal, format conversion, and indexing was performed with Picard (http://picard.sourceforge.net/). The Genome Analysis Toolkit was used to recalibrate base quality scores, perform local realignments around indels, and to call and filter the variants^15^^,^^16^. SnpEff was used to annotate gene and effect information for the variants^17^. Subject QC was performed using typical methods to check subject’s call rate, heterozygosity outliers, gender check, relatedness and population outliers. These QC checks were performed using Plink^18^ based on the SNVs that are in the intersection of the exome capture types used. VCFTools (http://vcftools.sourceforge.net/) was used to convert the SNV vcf file to Plink formatted genotypes. Population outlier checks also included subjects, populations and genotypes from the 1000 Genomes Project^19^, Phase1 Release v2.20101123. Individual genotypes were removed with genotype quality Phred-scores below 40. Omni-1M, Immunochip data and exome sequencing data were then merged with primary emphasis placed on the genotyping array data, only overwriting calls from arrays when genotyping data was missing. MaCH^11^ was used to determine phasing and the resulting files were converted into a variant call format (VCF) file using R script. An average of 286,824 SNPs with heterozygous genotypes were obtained per individual.

### eQTL discovery and replication

Prior to performing eQTL analyses, the Probabilistic Estimation of Expression Residuals^20^ (PEER) method was used to identify unknown factors affecting expression levels and so optimise eQTL discovery. PEER was run using default parameters with brain regions, age and gender accounted for as known covariates to generate 13 unknown factors (captured through the PEER axes), which were applied to RPKM normalised values to produce residuals. As would be expected, many of the 13 unknown factors were highly correlated with recognised covariates such as RIN and library batch (Supplementary Figure4). We also identified nine factors which were highly correlated with the specific brain region analysed. In fact, we found that expression of SLC6A3, which encodes the dopamine transporter (expressed with very high specificity in dopaminergic neurons of the ventral midbrain including the substantia nigra, see below), was highly correlated with the 10th PEER axis (p-value: 1.9 x 10 -26).

Variants within a ∓ 1Mb span of each normalised expression feature (i.e. gene, exon, exon-exon junction and unannotated intergenic region) were tested for cis-eQTL discovery using the R package MatrixEQTL^21^. Gender, age and the first three genetic principal component vectors were included as covariates within the linear model of marker genotype (imputed expected counts of minor allele) against normalized expression values. The Benjamini–Hochberg method was applied to calculate the false discovery rate (FDR) adjusting for the number of tests performed for each transcriptomic feature. Finally, we used a stepwise conditional analysis for each quantification type to detect independent variant effects targeting the same expression feature.

Three separate data sets were used to validate eQTL results: i) eQTL data reported by Ramasamy and colleagues^2^, which used an overlapping set of donor samples but a hybridisation-based microarray method for exon-level transcriptome quantification, ii) eQTL data generated by the GTEx consortium^22^, which was based on an independent sample set, but assayed similar brain regions and which used RNA sequencing to quantify the transcriptome, and iii) eQTL data relating to 373 lymphoblastoid cell lines generated by the Geuvadis consortium^23^, which used RNA sequencing for transcriptome quantification. Gene-level eQTLs (ge-eQTLs) identified within our study were declared as replicated when the significant SNP-gene pair was declared a hit with FDR < 5% in both our data set and the comparison data set.

### Characterisation of unannotated transcribed intergenic regions

We leveraged information from split reads (reads aligning to a genomic location with a gap or multiple gaps) and combined this with information on co-expression and physical proximity to known genes to characterise novel transcribed regions. In the first instance, the split read information for each sample was collected from the Tophat2^4^ junction output file, and overlaps between transcribed intergenic regions targeted by eQTLs and split reads were assessed using the GenomicRanges^24^ R package. If a transcribed region overlapped with a split read, then the same split read was screened for overlap with known genes. Only transcribed intergenic regions with split reads present in at least 4 separate samples were classed as having “high” evidence for being part of a known gene. In the absence of relevant split read data, the nearest gene (in genomic distance) from each transcribed intergenic region was selected using the GenomicRanges^24^ R package and the r² was calculated between the expression of the transcribed intergenic region and each exon of the nearest gene. Transcribed regions that had a maximum r² > 0.2 and were < 5Kb from the nearest gene were classed as having “moderate” evidence for being part of a known gene. Transcribed intergenic regions were classed as having “low” evidence for being part of a known gene when the r^2^ was < 0.2 or they were >5Kb from the nearest gene.

We also explored the possibility that unannotated transcribed regions could represent enhancer RNAs (eRNAs). We used the GeneHancer database v4.4^25^, which draws information from a wide range of sources (including ENCODE, the FANTOM5 atlas, the VISTA Enhance Browser, dbSUPER and EPDnew and UCNEbase) to define enhancer locations and then quantified the percentage overlap between eQTL target regions and enhancer regions. We adjusted for differences in the genomic size of eQTL target regions by calculating the percentage overlap per Mb of transcribed target sequence.

### Validation of unannotated transcribed intergenic regions in silico and by Sanger sequencing

We validated the transcription of intergenic regions targeted by i-eQTLs in silico using the data available on tissue-specific transcription generated by the GTEx consortium ([www.gtexportal.org](http://www.gtexportal.org)) and mapped through recount2^26^ The genome coordinates of all intergenic regions of interest were quantified from the transcription expression profiles generated with derfinder and available in recount2^26^. Counts were transformed to RPKMs and validation of transcription was considered when the region had an RPKM > 0.1 in at least 80% of samples in the tissue of interest.

We selected eight transcribed intergenic regions targeted by i-eQTLs for validation by Sanger sequencing. All regions were detected through analysis of putamen samples and we used a subset of RNA samples from the set of 111 used for RNA sequencing (**Supplementary Table 8**). In each case, reverse transcription was performed with 500 µg of total RNA using High Capacity cDNA RT Kit (Applied Biosystems) and random primers as per manufacturer’s instructions. PCR was performed using specific primers (**Supplementary Table 8**), all of which were designed to span predicted exon-exon junctions, and FastStart PCR Master (Roche). Amplification of the predicted band was confirmed by agarose gel electrophoresis and following confirmation, enzymatic clean-up of PCR products was performed using Exonuclease I (Thermo Scientific) and FastAP Thermosensitive Alkaline Phosphatase (Thermo Scientific). Sequencing was performed using the BigDye terminator kit (Applied Biosystems). All sequences were viewed using CodonCode Aligner (V. 6.0.2).

### Beta-hetereogeneity testing of eQTLs identified within and around a gene

A mixed model approach was used to identify heterogeneity in eQTL signal strength (beta coefficient or slope) within a gene. To test for beta heterogeneity between ge-eQTL and each other eQTL class (namely gi-eQTLs, e-eQTLs, ex-ex-eQTLs and relevant i-eQTLs) targeting the same gene, two models were fitted: 1) a non-heterogeneous (single slope) model with allele dosage as the main effect and two random effects, namely the index of the individuals and the index of the gene or exon or exon junction, and 2) a heterogeneous (multiple slope) model containing the same terms, but with the addition of a fixed-effect allele dosage × exon-or-exon-junction index interaction term. The lme4 R package was used to fit both models. A p-value for the likelihood ratio test comparing the two models was generated with the R function anova() and an FDR of < 5% was applied to assess significance.

### Identification of disease-relevant eQTLs

We used the STOPGAP database^27^ (accessed on 20^th^ of March 2018) to access and sub-classify loci identified through genome-wide association studies (GWAS). eQTL-GWAS overlap was checked using all eQTLs passing an FDR < 5%. The percentage overlap and enrichment was calculated based on the total number of eQTLs identified. Additionally, summary statistics were obtained from Parkinson’s disease (with the exclusion of data generated by 23andMe) and schizophrenia GWAS^28^^,^^29^. We applied *coloc*^30^ to colocalise Parkinson’s disease and schizophrenia loci with our eQTL signals. For each locus with a GWAS p-value of < 1 x 10^-5^ we examined all SNPs available in both data sets within 1Mb of the GWAS SNP of interest, and ran *coloc* with default parameters and priors. We called the signals colocalised when Coloc H3+H4 > or = 0.8 and H3/H4 > or = 2.

### ASE signal discovery and replication

The ASE discovery pipeline was motivated by Rozowsky and colleagues^31^ and involved the creation of parent haploids using phased variant data to reduce the impact of mapping biases on ASE identification. Each individual’s haploid genome was constructed using genotype data in the relevant VCF file, as parental information was unavailable. Deviations from the reference genome present in the VCF file were used to update the reference and create an artificial genome (two haploid genomes originating from each parent) using the Personal Genome Constructor tool vcf2diploid^31^ (version 0.2.4). The haploid genomes were arbitrarily referred to as "parent1" and "parent2". Trimmed fastq reads were aligned to individual haploid parent genomes using Tophat^4^ and Bowtie2^32^ (version 2.0.6), following a transcriptome guided approach, setting parameters to acquire the best alignment and allowing up to 2 mismatches. The alignments acquired for both haploid genomes , for each individual, were merged using the Suspenders tool (version 0.2.3), selecting the single best quality alignment. IGVTools^33^^,^^34^, version 2.3.18, was used to count the reads with one or the other hetSNP allele. Counts of both alleles, C1 and C2; alleles ordered alphabetically, were used to calculate the adjusted ratio $\left( C1+0.5 \right)/\left( C2+0.5 \right)$ to measure the ASE effect size. The use of +0.5 as an adjustment was motivated by the predicted reduction in the Taylor-Maclaurin bias estimator. Statistical tests for ASE were carried out via exact tests for departure from binomial expectation under the null hypothesis of $C1=C2$, using the binom.test() function in R. Tests were only conducted at heterozygous SNP sites where the total number of counts $\left( C1+C2 \right)$ exceeded 5 reads (“valid sites”). For each sample, the number of tests conducted was calculated and we converted the p-values into false discovery rates (FDR) using the Benjamini-Hochberg procedure. An FDR threshold of 5% was used to assess ASE significance in the valid sites. ASE signal validation was performed using ASE data generated from previously published lymphoblastoid cell line data ([Lappalainen](http://www.nature.com/nature/journal/v501/n7468/nature12531/metrics/blogs" \l "auth-1) et. al., 2013). After filtering for SNPs analysed in both data sets (n=54,214) we declared as validated those ASEs discovered in our brain data set that also had an FDR p-value of < 5% within the lymphoblastoid data set.

### Characterisation of ASE signals

Variant Effect Predictor^35^, SNPeff^17^ and SPIDEX^36^ were used to annotate all hetSNPs and to obtain a list of genes that had at least one significant hetSNP. For a subset of ASEs, namely those annotated as protein truncating variants within VEP (defined as stop gain, donor splice site and donor acceptor mutations) or common across individuals (defined as a ASEs identified in >= 10 individuals), we also investigated the impact of genotype on gene expression. The average expression of the exon or gene (following PEER axis correction) was calculated separately for individuals homozygous or heterozygous for the variant of interest or if possible across all three genotypes. We then performed t-tests or ANOVA as appropriate to identify departures from the null hypothesis of no association between expression and genotype.

### Investigation of the disease-relevance of ASEs using a bootstrapping approach

We investigated the enrichment of risk loci for Parkinson’s disease (with the exclusion of data generated by 23andMe)^29^ and schizophrenia^28^ amongst ASEs versus non-ASE sites by using a randomization approach similar to that described by Nicolae and colleagues^37^ to generate an empirical p-value for the significance of overlaps between risk SNPs and ASE sites, accounting for differences in read depth.

Let *C* be the set of SNPs that are in common between our set of hetSNPs (used for our ASE analyses) and the set of variants available from the GWAS summary statistics of the disease association study. Let *Ase* be the subset of SNPs in *C* that were declared to be ASE hits in our study, and let *NonAse* be the remaining subset of SNPs in *C* that were not declared to be ASE hits. For the randomization procedure, we repeated the following two steps 10^5^ times:

i) Randomly select SNPs with replacement from *NonAse* to create a SNP set with the same size and read depth distribution as that of *Ase*. This was done by placing the *NonAse* into bins based on their average read depth across samples and randomly selecting SNPs from the same to match the average read distribution of the *Ase*.

ii) Calculate the enrichment of GWAS signals within this SNP set by recording the GWAS p-values for each SNP within the set and computing the mean of the –log10 transformed values. Let r_i_ be the mean value for the i-th iteration.

The z-score for *Ase* (z_Ase_) was obtained using the mean and standard deviation of the r_i_ values, and was tested using the pnorm() function in R to obtain a p-value for the enrichment of GWAS risk SNPs amongst ASEs versus non-ASEs.

Using a similar approach, we also tested ASEs for evidence of enrichment of eQTLs and SNPs with effects on exon inclusion as predicted by SPIDEX.

### Assessment of brain cell type specificity for eQTL and ASE signals

We used the WGCNA R package^38^ to construct separate gene co-expression networks for substantia nigra and putamen, using as input for each network construction genes detected in > 70% of samples and with an RPKM of > 0.1. Following normalization using CQN^10^ and accounting for GC content, we used 13 PEER axes (see above) in addition to gender, age and genetic axes to perform covariate correction. The number of PEER axes was optimized to maximize the clustering of high quality cell markers^39^. We generated signed networks using a soft-thresholding power of 7 for substantia nigra and 8 for putamen networks, which approximated a scale-free topology. We obtained first-pass definitions of the modules via the dynamic Tree Cutting algorithm in WGCNA, and we then refined these modules by applying a k-means algorithm as described in our previous publication^40^. Modules were annotated in terms of cell-specific enrichments using the userListEnrichment R function implemented in the WGCNA R Package, combined with additional enrichment analysis on other marker gene sets^39^^,^^41-43^. Genes assigned to modules significantly enriched for brain-related cell type markers and with a module membership of > 0.3 were allocated a cell type “label” of neuron, microglia, astrocyte, oligodendrocyte and endothelial cell.

Next, for each eQTL targeting a known genic region, if the target gene was allocated to a cell then the related eQTL received the same cell type label. In the case of eQTLs targeting unannotated transcribed intergenic regions with high or moderate evidence for association to a known gene, the eQTL received a cell type label based on that known gene’s network position. For eQTLs targeting transcribed intergenic regions with low evidence for association with a known gene or which could not be classified, we assigned the target expression feature to a module (and by inference a cell type) based on its highest module membership (defined as the correlation of expression with the first principal component (‘eigengene’) of each module), provided the module membership was at least 0.3. Finally, for each eQTL class and each cell type (neuron, microglia, astrocyte, oligodendrocyte and endothelial cell) we applied a Fisher’s Exact test to test for enrichment of that cell type label amongst the genes associated to the eQTL class, relative to the genes not associated with that eQTL class but with module membership >0.3 to a relevant cell-type module. We followed a similar approach for ASEs and assessed ASE-containing genes for significant cell type enrichment.

### Partitioned heritability analysis of sites of allele-specific expression

GWAS summary statistics were obtained for schizophrenia^28^ and Parkinson’s disease (with the exclusion of data generated by 23andMe)^29^. Stratified LD score regression^42^, a method for partitioning SNP heritability across functional genomic annotations, was used to test for enrichment of schizophrenia and Parkinson’s disease heritability within ASE annotations. Region-specific annotations included putamen and substantia nigra (ASEs with FDR < 0.05 within a brain region were assigned to that region). In addition, an annotation (all) containing all ASEs irrespective of brain region with FDR < 0.05 was included. Annotations were added individually to the ‘baseline’ model of 53 annotations provided by Finucane et al., which comprises 24 different genome-wide annotations reflecting genetic architecture, such as conserved regions and histone marks. HapMap Project Phase 3 SNPs and 1000 Genome Project European population SNPs were used for the regression and LD reference panel, respectively. Only SNPs with minor allele frequency > 5% were used for heritability partitioning and the HLA region was excluded. We report the regression coefficient p-value, which tests whether the ASE annotations contribute significantly to SNP heritability after controlling for the effects of the ‘baseline’ model.

### Data availability

We have made data available in two primary formats. Using our web resource, http://braineacv2.inf.um.es/, users can access and visualise all forms of transcriptome quantification, eQTLs as well as gene co-expression networks (**Supplementary Figure 1**). The RNA-seq, whole exome sequencing and genotyping data can be accessed through the European Genome-phenome Archive numbers EGAS00001002113 and EGAS00001003065.
