## Supplementary figures and images for "Regulatory sites for known and novel splicing in human basal ganglia are enriched for disease-relevant information"

### Suppmentary Figure 1

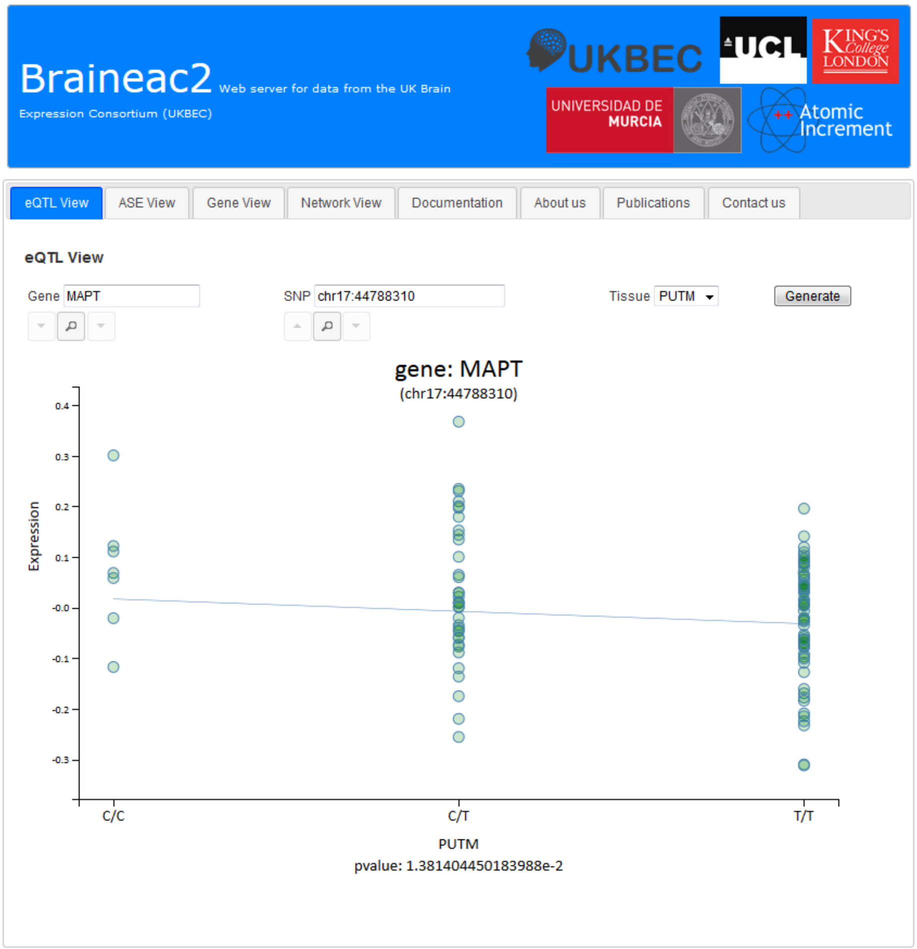

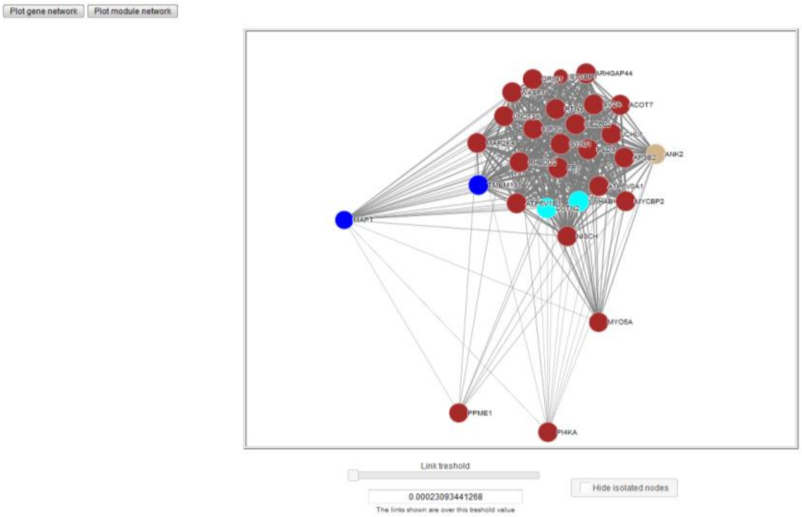

### Suppmentary Figure 2

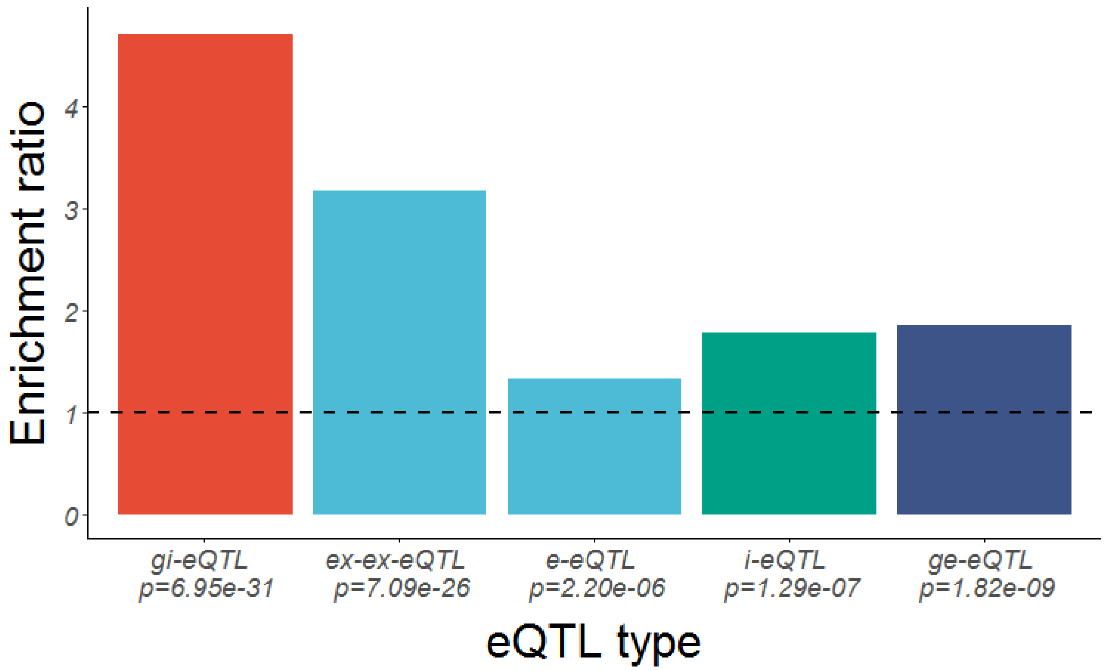

### Suppmentary Figure 3

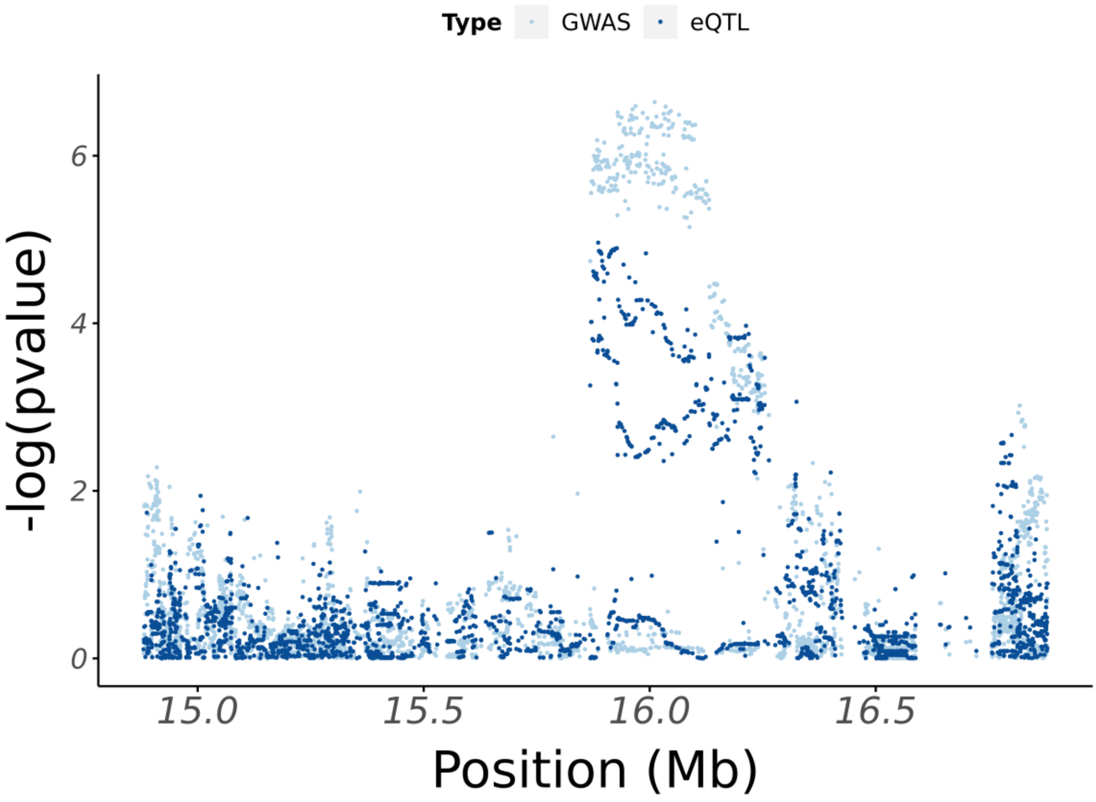

### Suppmentary Figure 4

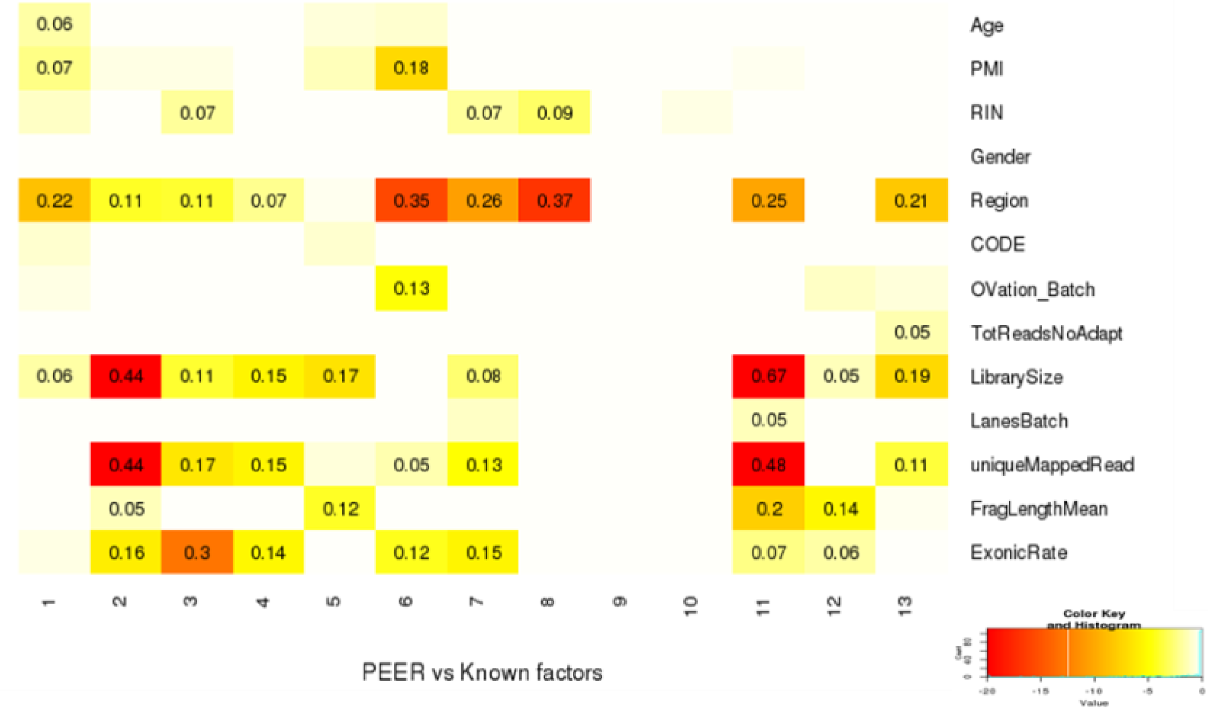
